# Plateau potentials are instructive signals for behavioral timescale synaptic plasticity in the neocortex

**DOI:** 10.1101/2025.11.07.687250

**Authors:** Courtney E. Yaeger, Raúl Mojica Soto-Albors, Weizhuo Liu, Anna Beltramini, Mark T. Harnett

**Affiliations:** McGovern Institute for Brain Research, MIT, Cambridge, MA, USA; Department of Brain and Cognitive Sciences, MIT, Cambridge, MA, USA

## Abstract

Learning occurs via the adjustment of synaptic weights across a variety of timescales. The mechanisms supporting these processes, from single-shot to iterative learning, are unclear. The prevailing model in the neocortex, spike-timing dependent plasticity (STDP), requires many pairings of precisely coordinated activity. This is difficult to reconcile with single-shot learning in behaving animals. In hippocampus, an alternative form of plasticity driven by plateau potentials (behavioral timescale synaptic plasticity; BTSP) alters synaptic weights in a single trial to reorganize spatial representations. Here we show that layer 5 pyramidal neurons (L5 PNs) of mouse primary visual cortex (V1) exhibit highly prevalent plateau potentials that drive single-shot changes in sensory representations. Spontaneously occurring and experimentally induced plateaus rapidly and persistently modified L5 PN responses to visual stimuli. In acute slices, plateau potentials drove synapse-specific plasticity with few repetitions across seconds-long pairing intervals. Our results demonstrate that plateau potentials in the neocortex rapidly reshape neuronal representations through BTSP. This instructive form of plasticity provides a mechanism for dynamic adjustment of neocortical synaptic weights to support rapid learning.

## INTRODUCTION

Many forms of synaptic plasticity have been described in the neocortex, yet it remains unclear how these contribute to learning in behaving animals. The predominant model, Hebbian spike-timing dependent plasticity (STDP), requires dozens of precisely timed pre- and post-synaptic pairings^1–8^. This is at odds with both the highly variable nature of neocortical spiking and the ability of animals to learn from single or few experiences^9–11^. Even modified versions of STDP^12–18^ have difficulty explaining how behaviorally relevant synapses are rapidly modified. In addition, the repetition requirement of STDP precludes its involvement in single-shot learning. Thus, rapid learning likely requires plasticity mechanisms beyond conventional STDP.

Behavioral timescale synaptic plasticity (BTSP) is a one-shot learning rule first discovered in hippocampus in which a single plateau potential instructs place field formation^19–22^. Plateau potentials are prolonged somatic depolarizations that span tens to hundreds of milliseconds and are crowned by high frequency action potential (AP) firing^19,23,24^. BTSP allows the hippocampal network to rapidly produce spatial representations that reflect behaviorally salient features^25–30^. This instructive plasticity mechanism may underlie more general forms of learning if expressed in other brain regions. Whether the neocortex uses BTSP to rapidly reshape representations remains unknown.

## RESULTS

### Plateau potentials are highly prevalent in neocortical L5 PNs in behaving mice

Plateau potentials are commonly observed in hippocampal pyramidal neurons in vivo and have been well characterized using whole-cell patch-clamp recordings^19,22,24,31,32^. Observation of these events in the neocortex, however, is limited to a few reports. Plateaus have not been reported in any of the extensive patch-clamp studies conducted in layer 2/3 of behaving animals, aside from small subthreshold depolarizations mediated by NMDARs^33,34^. An isolated observation of plateau-like waveforms in the awake neocortex lacked laminar identification^35^, and one group reported plateau potentials in layer 5 (L5) PNs of the primary visual cortex (V1) under anesthesia^36,37^.

To characterize plateau potential properties and prevalence in behaving mice, we performed whole-cell recordings in V1 L5 PNs in awake mice viewing naturalistic visual stimuli. Recordings were made from awake head-fixed mice that were free to locomote on a treadmill while passively viewing movie clips (Fig. 1a). These recordings revealed numerous instances of high-frequency action potential firing crowning large, sustained depolarizations (Fig. 1b), surprisingly similar to the plateau potentials previously defined in the awake hippocampus. Using similar criteria to those used in hippocampus^19^, we defined neocortical plateaus as prolonged depolarizations (≥10 ms) during which the smoothed membrane potential (V_s_) exceeded -35 mV and two or more action potentials were present (Fig. 1c, Extended Data Fig. 1). We identified plateau potentials in 45 out of 83 L5 PNs from 63 mice. In those neurons, we recorded a total of 4097 plateau events, with a mean duration of 48 ± 1.5 ms (IQR 19-51 ms, Fig. 1d). Plateau events in L5 occurred at strikingly high frequencies (0.16 ± 0.03 Hz) but were similar to those seen in CA1 neurons when normalized to firing rate (0.97 ± 0.17 plateaus per 100 spikes, Fig. 1e)^19^. Post-hoc immunohistochemistry revealed that plateaus were present in both extratelencephalic (ET) and intratelencephalic (IT) L5 PNs (Extended Data Fig. 2). Pyramidal neurons in layers 2/3 (n = 11) and 4 (n = 11) did not exhibit plateaus according to our criteria (Fig. 1f, Extended Data Fig. 1). These findings establish the plateau potential as a widespread feature of L5 PNs in awake mice and a potential mechanism to induce rapid changes in sensory output.

**Figure 1.**
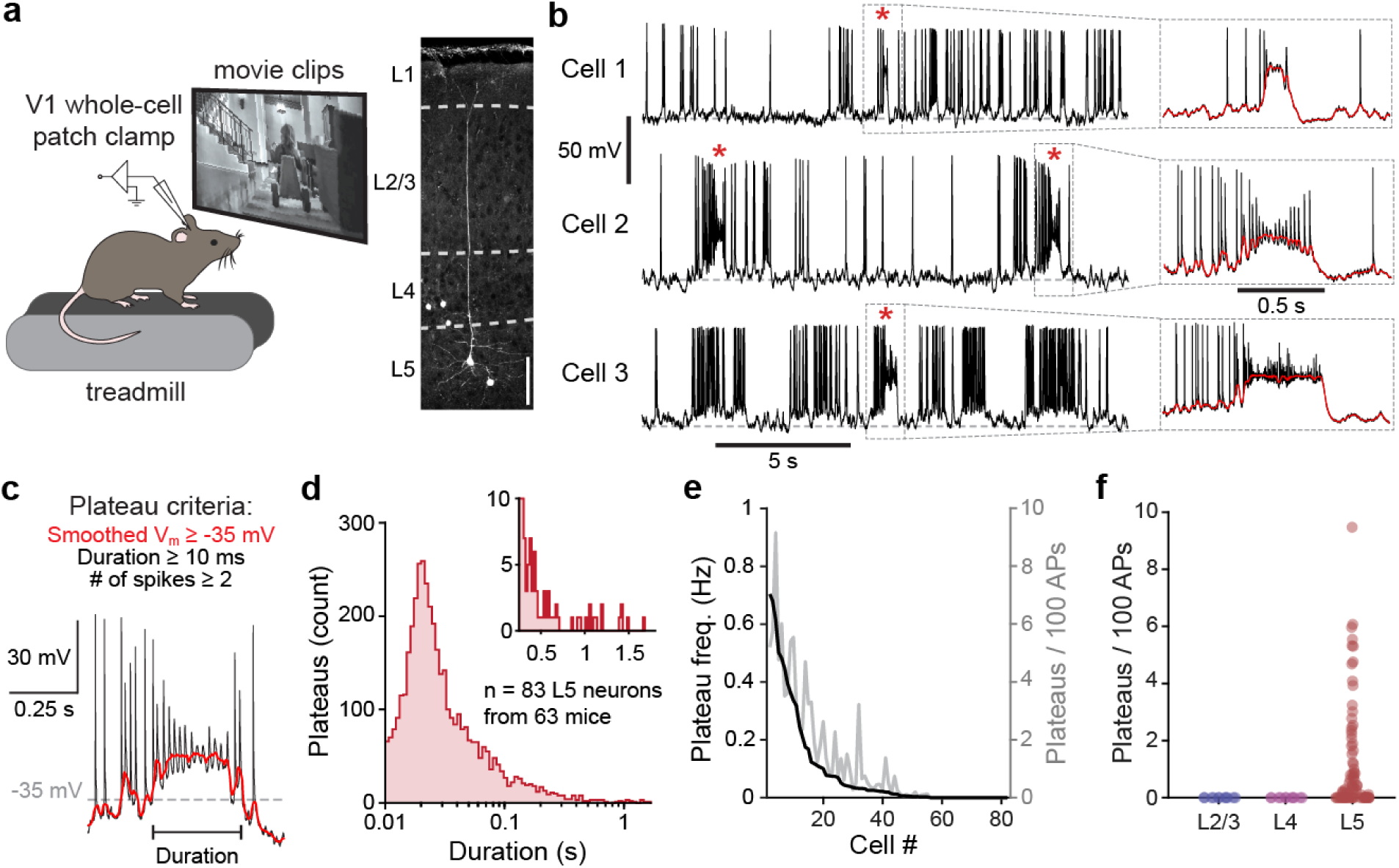
Plateau potentials are highly prevalent in L5 pyramidal neurons. **a**) Experimental setup (left) and confocal reconstruction of a L5 PN filled with biocytin (right). Scale bar: 100 µm. **b**) Left: representative intracellular V_m_ traces from three L5 PNs, with plateau potentials indicated by red asterisks. Right: expanded plateau potentials from the corresponding traces on the left, with smoothed V_m_ in red. **c**) Plateau potential criteria and associated example trace. **d**) Duration distribution for all plateau potentials detected in L5 PNs during movie presentations (n = 4097 events from 83 cells across 63 mice); inset shows distribution tail over 250 ms. **e**) Plateau frequency in Hz and events per 100 APs for each L5 PN. **f**) Plateaus per 100 APs across PNs in different cortical layers (L2/3 n = 11 cells, L4 n = 11 cells, L5 n = 113 cells).

Plateau potentials in hippocampus occur spontaneously in both silent cells and those that already exhibit place field firing^19,21,22^. However, some systematic factor must drive plateaus to ultimately produce behaviorally-relevant representations. To determine whether neocortical plateau potential generation is a function of sensory input, behavior, or internal state, we recorded locomotion, orofacial movements, and pupil area during movie clip and static grating presentations (Extended Data Fig. 3, see Methods). Plateau rate did not differ between stimuli and intertrial intervals (Extended Data Fig. 3b), nor between stationary and locomoting periods, quiet and whisking epochs, or pupil dilation and constriction states (Extended Data Fig. 3c-d). However, when behavioral variables were aggregated into an overall activity metric, we observed significant plateau rate elevation during active epochs (Extended Data Fig. 3c-d). These results suggest that the occurrence of plateau potentials is not directly driven by visual stimulation or behavioral variables but may be weakly modulated by general arousal.

### Neocortical plateau potentials rapidly modify neuronal representations

Plateau potentials drive rapid changes in feature selectivity in the hippocampus, where a single plateau potential can transform silent cells into place cells or shift place field firing from one location to another^19–22,26,27,32,38^. V1 L5 PNs exhibit selective responses to sensory stimuli, including movies^39–41^, raising the question of whether plateau potentials could alter these representations in single neurons. A subset of L5 PNs exhibited a sudden change in response to movie presentation after a long spontaneous plateau potential (n = 7, Fig. 2a-c). Subsequent trials had a significant and persistent increase in V_s_ and a corresponding elevation of firing rate (ΔV_s_ = 5.68 ± 0.46 mV; ΔHz = 6.53 ± 1.05 Hz; Fig. 2d-f, Extended Data Fig. 4). Significant changes in V_s_ were specifically associated with plateaus longer than 200 ms (>200 ms: 1.10 ± 0.28 s, 7 plateaus from 7 cells; <200 ms: 0.04 ± 0.01 s, n = 16 plateaus from 5 cells; Fig. 2g), and plateau duration was correlated with the resulting magnitude of change in V_s_ (Fig. 2h). This is consistent with findings in hippocampus, where spontaneous plateaus that induce place fields are approximately three times longer than plateaus in existing place fields^19,21,22^. These results demonstrate that long-duration plateau potentials in V1 L5 PNs can rapidly modify neuronal tuning, similar to their role in hippocampal place field formation.

**Figure 2.**
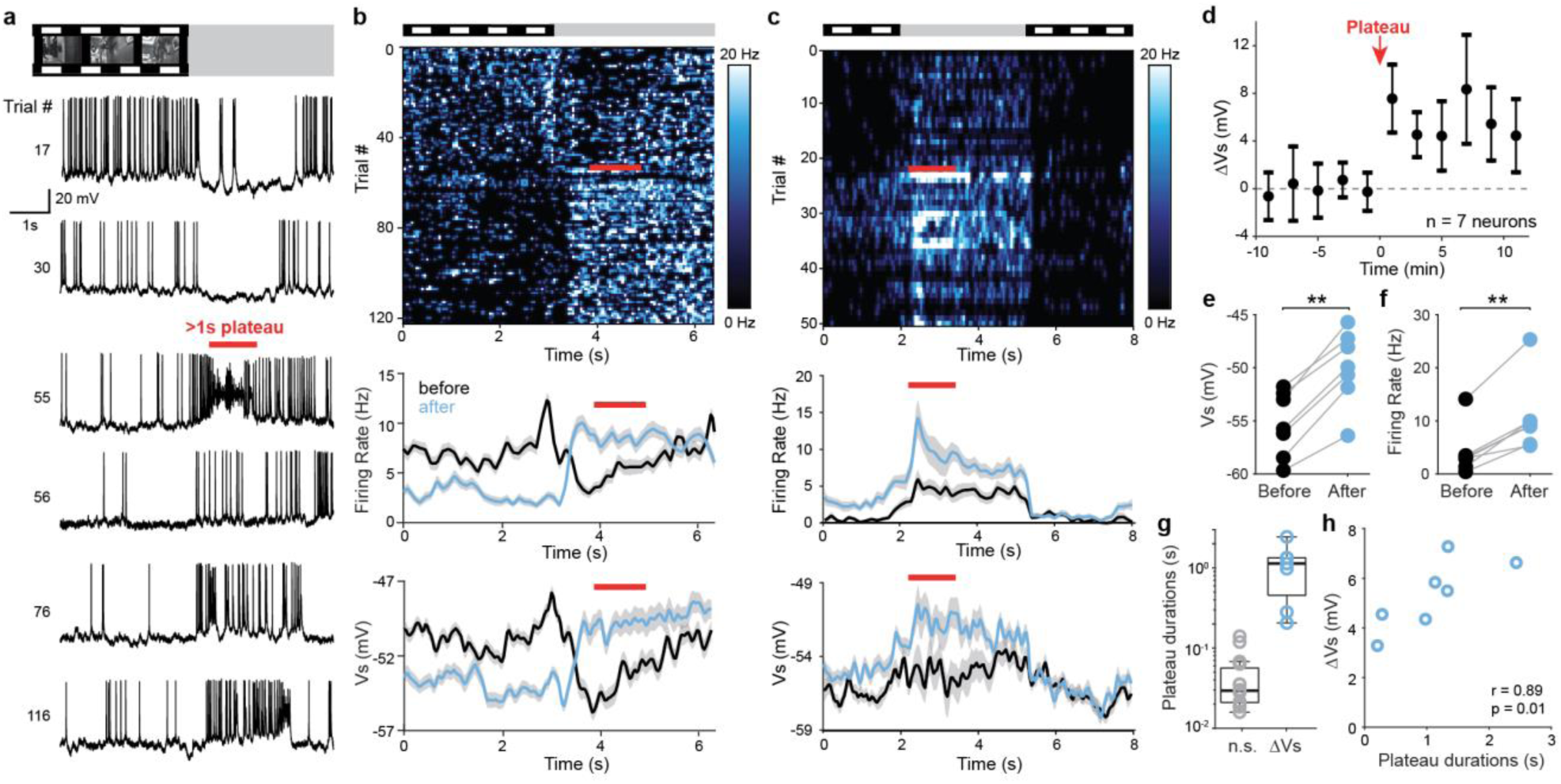
Spontaneously occurring plateau potentials are associated with rapid modification of neuronal responses in awake mice. **a)** Representative intracellular V_m_ from a V1 L5 PN for select trials before, during, and following a long plateau potential (trial 55, red bar). Schematic at top indicates timing of presented movie and interleaved grey screen. **b)** Top: firing rate for each trial for the cell in a, as a function of time. Middle and bottom: average firing rate and average V_s_, respectively, for trials before (black) and after (blue) the plateau occurred. Red bar indicates the start and duration of the long duration plateau in trial 55. **c)** As in b, for a second representative V1 L5 PN with a long duration plateau potential**. d)** Average ΔV_s_ across time for all cells with long duration (>200 ms) plateau potentials. **e)** Average V_s_ for trials before and after spontaneous long duration plateaus for the cells in d. Wilcoxon signed-rank test, p = 0.01. **f)** As in e, but for firing rate. Wilcoxon signed-rank test, p = 0.01. **g)** Plateau durations associated with a lack of significant ΔV_s_ (grey, n=16 plateaus from 5 cells) versus those with significant ΔV_s_ (blue, n=7 plateaus from 7 cells). **h)** Average ΔV_s_ (after-before) as a function of plateau duration, for V_s_ changes reaching significance criteria (n = 7 plateaus, 7 cells). Spearman’s correlation and p-value shown. Data: mean ± SEM. Boxplots indicate the 25th and 75th quartiles, with whiskers extending to ±2.7σ.

If plateau potentials serve as instructive signals for rapid synaptic plasticity in L5 PNs, experimental induction should also modify neuronal responses. We therefore paired prolonged current injections with movie presentations to mimic plateau potentials (Fig. 3a). We injected a single second-long pulse (800 pA) in 11 neurons and 200-500 ms pulses across multiple trials in 3 other neurons (5-20 pairings, Extended Data Fig. 5). These current injection durations spanned the range of the spontaneous plateaus we observed previously to drive significant changes in membrane potential (Fig. 2h). In each experiment, the current injection occurred at a randomly chosen time during the paired movie. Experimental plateau induction produced a rapid increase in V_s_ and increased spiking only in the paired movie around the induction time and in the trials immediately following current injection (Fig. 3b). These changes in V_s_ persisted throughout the recording (ΔV_s_ = 4.09 ± 0.46 mV, n = 14 neurons; Fig. 3d). A second unpaired movie was pseudorandomly interleaved with the paired movie and served as a control (Fig. 3c). The unpaired movie showed no significant change in V_s_ or firing rate at the timepoint of peak ΔV_s_ in the paired movie (ΔV_s_ = 0.63 ± 0.40 mV, firing rate ratio = 0.92 ± 0.15 Hz; Fig 3c-g). These results demonstrate that plateau-like depolarizations are sufficient to increase input efficacy and enhance spiking output.

**Figure 3.**
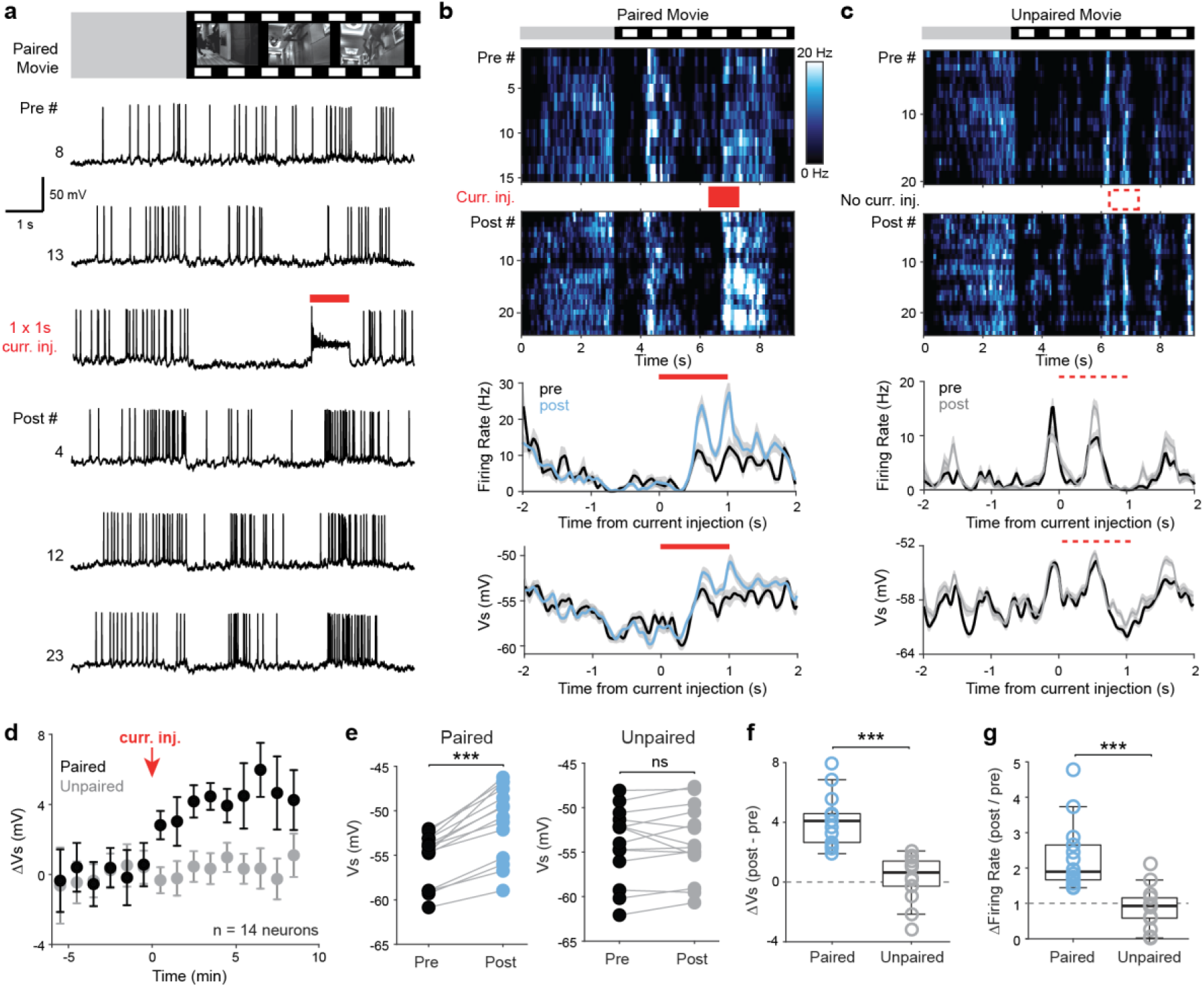
Induced plateau potentials enhance feature selectivity. **a)** Representative intracellular V_m_ for select trials before, during, and following a single 1-second current injection to induce a plateau potential (red bar). Schematic at top indicates the end of the grey screen intertrial interval and the start of the movie presentation. **b)** Top: firing rate for pre- and post-induction trials for the cell shown in a as a function of time. Middle and bottom: average firing rate and average V_s_, respectively, before and after induction. Red bar indicates the start and duration of the current injection. **c)** As in b, for the same cell’s responses to a second movie, which was not paired with a current injection. Dashed line indicates the start and duration of the current injection in the paired movie. **d)** Change in V_s_ as a function of time for paired and unpaired movies pre- and post-induction. **e)** Left: average V_s_ pre- and post-induction in paired movies. Wilcoxon signed-rank test, p = 1.22E-4. Right: average V_s_ pre- and post-induction in unpaired movies. Wilcoxon signed-rank test, p = 0.42. **f)** Average change in V_s_ for paired and unpaired movies. Wilcoxon signed-rank test, p = 1.22E-4. **g)** Average change in firing rate for paired and unpaired movies. Wilcoxon signed-rank test, p = 1.22E-4. Data are mean ± SEM. Boxplots indicate the 25th and 75th quartiles, with whiskers extending to ±2.7σ.

Plateaus delivered at randomly chosen timepoints in the movies drove reliable changes in V_s_, but the magnitude of these changes was variable and correlated with the preinduction firing rate of the same epoch (Extended Data Fig. 5). This variability suggests that plateau-mediated plasticity may be limited to specific inputs. Orientation selectivity in V1 is primarily driven by thalamocortical inputs^42–45^, which cannot be potentiated in L5 PNs of adult mice^46^. Therefore, we conducted a set of experiments in which we identified each cell’s orientation preference to static gratings and then paired a prolonged current injection with a randomly selected non-orthogonal, non-preferred angle (300 ms, ≥600 pA, 10 repetitions; Extended Data Fig. 6). Although we did observe a slight but significant reduction in the responses to the preferred orientation (ΔHz = - 1.61 ± 0.55 Hz, n = 14), this paradigm did not increase responses at paired orientations (ΔHz = 0.09 ± 0.50).

To distinguish whether the depolarizing envelope or the high-frequency AP firing of plateaus drove rapid changes in neuronal responses, we paired movies with a second-long train of high-frequency action potentials without sustained depolarization (60 Hz, 2 ms current injections for 1 s; Extended Data Fig. 7). No significant changes in V_s_ were detected (ΔV_s_ = -0.44 ± 0.40 mV, n = 6 neurons), demonstrating that the depolarizing envelope of plateau potentials, rather than high-frequency firing, drives changes in movie responses.

### Neocortical plateau potentials support a behavioral timescale synaptic plasticity rule

Plateau potentials evoke rapid, robust single-shot synaptic plasticity in CA1 and CA3^19,22,26,27,38^, potentiating inputs according to a seconds-long timing rule^20,21^. We tested if plateau potentials could induce plasticity at similar timescales in V1 L5 PNs with whole-cell patch-clamp recordings in acute slices from adult mice. Local electrical microstimulation was used to produce EPSPs (Fig. 4a). Following a ten-minute baseline, a train of 10-20 Hz EPSPs was paired with a somatic current injection (300 ms, 800 pA) at a set interval (-2 s to +0.5 s), mimicking L5 synaptic input and a long plateau potential, respectively. This pairing was repeated only 5 times with 15 seconds between epochs and then monitored for at least 30 minutes after induction, as has been previously done in CA1^20^. Overlapping the presynaptic stimulus train with the current injection (“simultaneous”, t = 0) evoked robust plasticity (normalized peak EPSP postinduction, 1.66 ± 0.13, n = 8 neurons, Fig. 4a,b, Extended Data Fig. 8). We observed substantial potentiation at non-overlapping timepoints, with pre-before-post pairings potentiating inputs up to a -2 s interval, and post-before-pre pairing potentiating at the +0.5 s interval (Fig. 4e, Extended Data Fig. 8).

**Figure 4.**
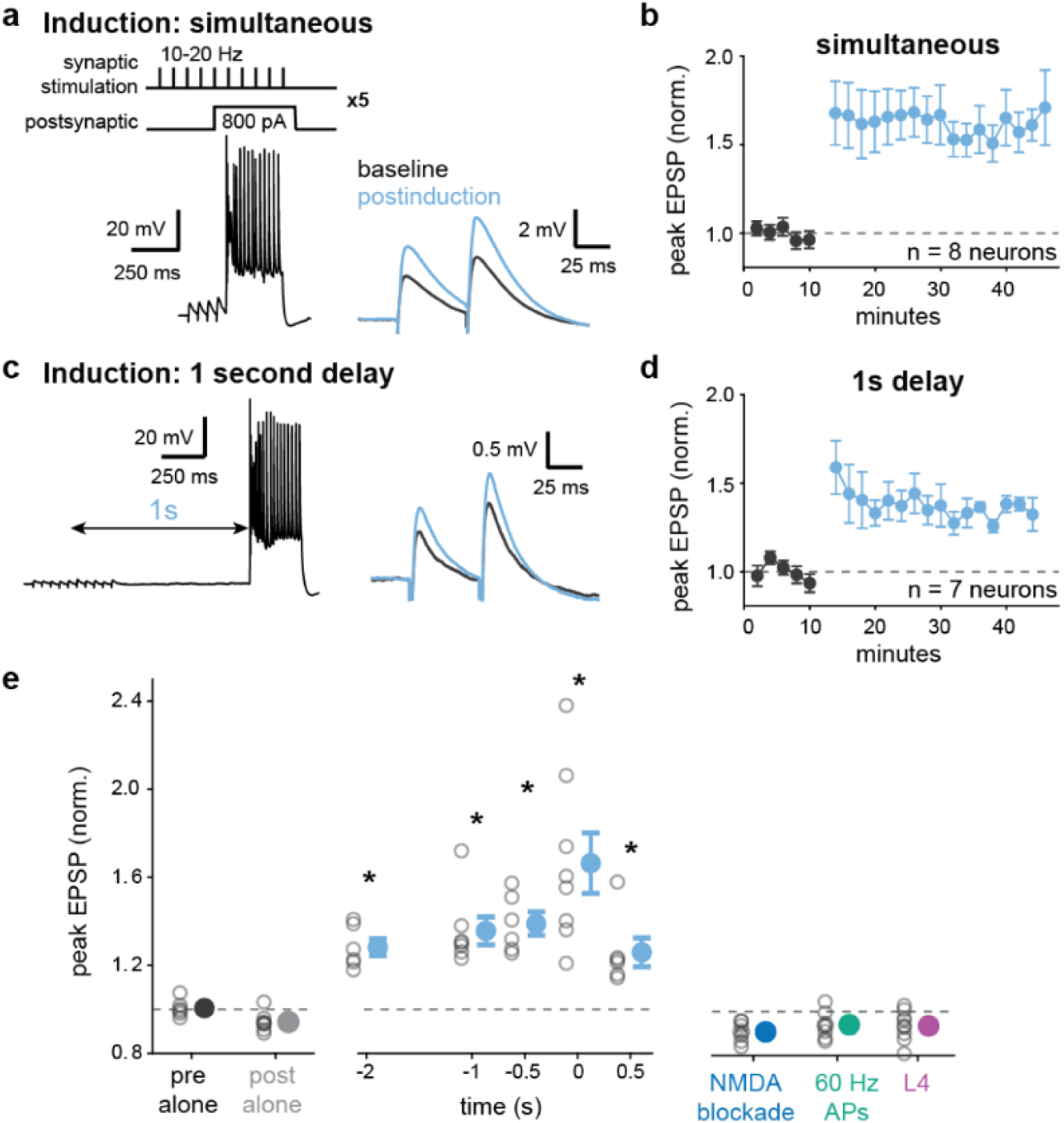
Plateau potentials drive cortical synaptic potentiation at behavioral timescales. **a)** Left: schematic (top) and representative voltage trace (bottom) of the simultaneous (t = 0) induction paradigm, in which a train of EPSPs driven by electrical stimulation is paired with a somatic current injection 5 times. Note the overlap of the current injection with the middle of the EPSP train. Right: average paired pulse response during the baseline period (black) and the average post-induction response (blue). **b)** Normalized EPSP amplitude for a population of V1 L5 PNs that received the simultaneous induction paradigm. Induction occurs after 10 minutes. **c)** As in a, but for an induction protocol where the current injection occurs at a 1-second delay from the EPSP train. **d)** EPSP amplitudes before and following the 1-second delay induction paradigm. **e)** Mean post-induction changes in EPSPs for each induction timing (middle) as well as unpaired controls (left) and perturbations (right). Open circles are individual neurons; filled circles are means; bars are ± SEM.

We next tested the key mechanistic components of this rapid, long-timescale neocortical synaptic plasticity. Plasticity required coordinated pre-post pairing: presynaptic or postsynaptic activity alone did not produce any potentiation (Fig. 4e, Extended Data Fig. 8). Pairing-induced potentiation (at t = 0) was abolished by blockade of NMDA receptors (Fig. 4e, Extended Data Fig. 8). Potentiation also required prolonged membrane potential depolarization: when single action potentials were driven at a comparable frequency to those observed during the step current injection (60 Hz), synaptic strengthening did not occur (Fig. 4e, Extended Data Fig. 8). Additionally, synapses in the layer 4 apical oblique dendritic domain did not potentiate, likely due to the elevated AMPA-to-NMDA ratio found in these thalamorecipient dendrites^46^ (Fig. 4e, Extended Data Fig. 8). Our findings collectively demonstrate that long plateau depolarizations are instructive for a behavioral timescale plasticity rule in neocortical L5 PNs.

## DISCUSSION

We demonstrate that long plateau potentials in V1 L5 PNs drive single-shot changes in sensory responses. Our data is consistent with a model in which plateau potentials serve as instructive signals for a form of plasticity that spans seconds and does not require repeated, persistent co-activation of neurons^47^. Neocortical BTSP therefore represents a novel mechanism for one-shot learning.

Plateau potentials were surprisingly common in L5 PNs: we observed them in 45 out of 83 neurons. These plateaus occurred at 0.16 Hz, a rate similar to those in CA1 when normalized by firing rate (0.97 ± 0.17 plateaus per 100 APs)^19^. Only spontaneous plateaus with long durations (>200 ms) drove rapid selectivity changes, congruent with the duration of plateaus that evoke place fields in hippocampus^19,21,22^. Long plateaus sufficient for rapid plasticity are likely driven by convergent inputs, neuromodulation, disinhibition, or some combination of these factors^47–49^. The majority of events were short (∼48 ms) and did not induce plasticity, as is the case in hippocampus^19^. The role of shorter plateaus is unclear; they may evoke levels of potentiation undetectable with our approach or serve a different function entirely, such as gain modulation or multiplexing^19,50–54^. Dendritic electrogenesis may be involved in neocortical plateau generation^55–58^, as it is in CA1^19^. In awake animals, active dendritic events are also associated with large calcium transients and may carry task-specific information^59–64^ which could serve as instructive signals^65,66^.

We observed plateaus in both ET and IT L5 PNs, but not in L2/3 and L4 neurons. It remains an open question whether one-shot plasticity mechanisms operate in other layers, potentially using alternative waveforms such as high-frequency burst firing or local NMDA spikes^33,34,67–70^.

Our data indicate that BTSP in L5 PNs is limited to specific inputs. Plateaus failed to alter orientation selectivity, which is largely driven by dLGN input^42–45^. dLGN inputs on L5 PNs, which synapse at proximal apical dendrites, do not potentiate with either BTSP or STDP paradigms in vitro^46^ (Fig. 4). Congruent with this, we observed variable magnitudes of potentiation for induction across different movie frames. Together, these results suggest that plateau-mediated plasticity in V1 potentiates inputs other than the feedforward pathway. This architecture may enable L5 PNs to acquire context-dependent responses without overwriting the basic feature selectivity conveyed by primary thalamic inputs. This is congruent with previous work on single-neuron mechanisms for balancing flexibility and stability in learning systems^46,71–74^.

These results demonstrate that the neocortex possesses a mechanism for instructive plasticity capable of reshaping representations from single experiences. Our discovery supports the proposition that BTSP enhances behaviorally-relevant sensory, motor, and/or cognitive representations beyond hippocampus^47^. The differences we observed between neocortical and hippocampal BTSP may reflect distinct learning algorithms across brain regions^20,75–77^. Our data are consistent with biologically-inspired models of supervised learning in hierarchical networks^52,65,78^, in which AP bursts associated with plateau potentials enable propagation of instructive signals through feedback connections. Instructive plasticity via plateau potentials may therefore be a general principle for learning in hierarchical systems.

## METHODS

### Surgery and animal handling

All animal procedures were carried out in accordance with the US National Institutes of Health and the Massachusetts Institute of Technology Committee on Animal Care guidelines. Male and female C57BL/6 mice were housed in pairs in a reversed 12-hour light/dark cycle room with ad libitum food and water access. The room was kept at 20-22.5 °C and 30-70% humidity.

Mice were implanted with a headplate at 8-10 weeks of age. They were initially anesthetized with 3-5% isoflurane and subsequently maintained at 1.5-2% isoflurane throughout the surgical procedure. Briefly, the skull was exposed and cleared of connective tissue. The medial (1.5 mm) and lateral bounds (3.5 mm) of primary visual cortex (V1) were marked on the cranium using a permanent marker for future reference. Light pressure was then applied to the marked skull plate and sutures were fixed in place with cyanoacrylate glue. Subsequently, the headplate was attached using cyanoacrylate glue and cemented with clear Metabond (Parkell Inc), taking care to fully cover the titanium headplate and electrically isolate it from the skull.

Following implantation, mice were allowed to recover in their home cage for a minimum of 3 days before initiating habituation. All habituation was performed with minimal light to avoid light cycle disruption. Animals spent 1-2 days acclimating exclusively to experimenter handling. Once comfortable, mice were head-fixed on a flat polyester treadmill for the following 2-6 days for increasingly longer durations, ranging from 5-60 minutes. During later sessions, animals were habituated to the presence of rig and screen lights as well as experimenter movements over the craniotomy, including the placement of a physical barrier posterior to the craniotomy and over the ears to prevent the animal from disturbing the recording site. This was done to simulate experimental conditions to ensure animals were calm and stress-free on the recording day.

After successful habituation (2 or more weeks post-implantation, at 10+ weeks of age), a short surgery was performed on the day of recording. Mice were lightly anesthetized with 1.5-2% isoflurane. To prevent fur from invading the recording site, the animals’ neck and upper back hair was removed. A craniotomy (⌀ ∼0.3 mm) was performed within the previously stereotactically marked bounds, followed by a durotomy for unimpeded access to the brain. To seal and protect the exposed brain during recovery, 1.5% low-melting point agarose (ThermoFisher Scientific) was applied and a ⌀ 3 mm window was nestled within the agarose. Once set, a thick layer of Kwik-Cast (World Precision Instruments, USA) was applied to prevent the agarose, and thus the brain, from drying. Animals were allowed to recover for at least 60 minutes on a heating pad in their home cage before being transferred to the experiment rig. To reduce electrostatic build-up, the treadmill was treated with anti-static spray (ACL Staticide) before animals were head-fixed.

### In vivo electrophysiology

Whole-cell borosilicate patch electrodes (5-8 MΩ; World Precision Instruments) were filled with a solution containing (in mM): 134 potassium gluconate, 6 KCl, 10 HEPES, 4 NaCl, 4 Mg_2_ATP, 0.3 NaGTP, and 14 phosphocreatine di(tris). In a subset of experiments, biocytin was added to the solution at a concentration of 0.08-0.12% for histological reconstruction of patched cells. Patch electrodes were mounted on a micromanipulator (LN Mini, Luigs and Neumann) with an azimuth of 46-48° to prevent damage to apical dendritic arbors. Immediately before starting an experiment, the agarose plug was removed and replaced with cortex buffer, an isotonic solution containing (in mM): 125 NaCl, 5 KCl, 10 glucose, 10 HEPES, 2 CaCl_2_, and 2 MgCl_2_. The manipulator was zeroed when the pipette contacted the brain surface, as detected by a stereotypical deflection in the test pulse. The electrode was then lowered through the cortex at 2 µm/s with 55-65 kPa of pressure to prevent clogging. Upon reaching 100 µm, the pressure was steadily lowered to 15-20 kPa; upon reaching the top of the desired layer (150 µm, 400 µm, and 650 µm for layer 2/3, layer 4 and layer 5, respectively), the pressure was further reduced to 1.8-2.5 kPa to search for cells. Putative cells were identified by reproducible increases in electrode resistance. Following the formation of a gigaohm seal, capacitive currents were neutralized before break-in. After successfully achieving a whole-cell configuration, the amplifier was switched to current clamp for the duration of the experiment. Bridge balance was initially corrected upon break-in and subsequently monitored and corrected prior to the current injections during input-output or plasticity experiments. All data was acquired on a BVC-700A amplifier (Dagan Corporation) and low-pass filtered at 10 KHz, collected on a BNC-2110 terminal block (National Instruments), digitized at 20 KHz using a PCIe-6374 (National Instruments), and recorded on a modified version of WaveSurfer 0.986 (Howard Hughes Medical Institute) that allowed synchronous data acquisition from four analog channels. The liquid junction potential was not corrected.

### Visual stimulation

Presentation of visual stimuli was performed at 60 Hz using custom scripts written in MATLAB (Psychtoolbox-3^79^) and Python (PsychoPy^80^), and displayed on a 27” in-plane switching (IPS) liquid crystal display (LCD) monitor (Dell G2724D) placed 14-20 cm away from the mouse on the contralateral visual field.

#### Movie clips

Movie presentation consisted of 6 or 10 seconds of continuous scenes with a 3-5 second intertrial interval gray screen. Publicly available clips from the following films were used: Inception (2010), Gravity (2013), The Shining (1980), Fantastic Mr. Fox (2009), The Revenant (2015), Birdman (2014), The Protector (1985), and Touch of Evil (1958). Movies were contrast normalized and presented in greyscale.

#### Static gratings

Full-field static gratings were presented for 0.2 s and had varying intertrial interval gray screen durations (0.1, 0.5 or 1 s). The presented gratings consisted of 8 different orientations (0 -180°, 22.5° intervals) at 0.04 cycles/deg in pseudo-random order.

### Behavioral set-up and videography

Mice were head-fixed on top of a flat linear polyester treadmill measuring 28.5 cm in length and were free to walk at will. Treadmill movement was monitored using a quadrature optical encoder (Broadcom Ltd.) and converted to an analog signal using a Teensy 3.2 or a Teensy 4.0 (PJRC), collected on a BNC-2110 terminal block, digitized at 20 kHz using a PCIe-6374 (National Instruments), and recorded simultaneously with electrophysiological data on WaveSurfer 0.986. To track pupil dilation and orofacial movements, mice were illuminated with an infrared light and monitored using a Teledyne DALSA Nano-1280 GigE camera. Frame acquisition was triggered using the MATLAB Image Acquisition Toolbox at 60-62 frames per second. Frame synchronization was achieved by generating a TTL pulse each time a new frame was acquired.

### In vivo intracellular voltage data processing

Intracellular voltage was filtered post-hoc using a zero-phase 5 kHz second-order low-pass digital filter (Butterworth) to remove high-frequency noise. Any remaining outliers were detected, removed, and linearly interpolated using a z-score threshold of -7. To correct for baseline drift, voltage traces were split into discrete 10 second windows. For each window, the average deviation of AP thresholds at or below the fifth percentile (most hyperpolarized 5%) from an expected value of -42 mV was added to the trace. To quantify the average baseline V_m_, APs were removed by deleting all points from 0.25 ms before a threshold value (dV/dt ≥ 50 V/s) to 3.4x AP half widths after, and the missing values were linearly interpolated. The resulting traces were used to create histograms with a bin size of 0.1 mV, and each cell’s baseline V_m_ was defined as the mode of the distribution. For all analyses, we excluded neurons that met any of the following criteria: (1) total recording duration < 10 minutes, (2) baseline V_m_ > -45, and (3) high input resistance (Ri > 150 MΩ). Additionally, we excluded putative interneurons as identified by either short spike half-widths (< 0.5 ms) or stark spike afterhyperpolarization.

### Plateau potential characterization

To identify plateau potentials, we smoothed the AP-removed voltage using a 20 ms boxcar moving average. Any event that exceeded a threshold of -35 mV was labeled as a putative plateau. In line with previous literature^19^, this threshold was determined as an estimate of where smoothed V_m_ distributions deviated from a bimodal Gaussian fit. Plateau amplitude was calculated as the difference between plateau peak and baseline V_m_. Plateau duration was defined as the time difference between the positive- and negative-going threshold crossings. Putative plateaus that were shorter than 10 ms or had less than two spikes were eliminated. For plateau spiking analyses, spikes occurring within 25 ms of a plateaus’ rising and falling edges were labeled as plateau APs. Spike amplitude adaptation ratio was determined by dividing the minimum AP amplitude by the maximum AP amplitude within a plateau. Spike half widths were calculated using the MATLAB function findpeaks, and all widths were normalized to the mean half width of the first spike in a plateau. Plateau prevalence was calculated from the total number of plateaus divided by the total number of APs and multiplied by 100. Neurons were classified as exhibiting plateau potentials if they displayed at least three events during the recording session (mean ± S.E.M. plateaus per cell, L2/3: 0.09 ± 0.09, L4: 0.46 ± 0.31, L5: 58.08 ± 13.83).

### Behavioral correlates

Pupil area and whisking behaviors were determined offline using FaceMap^81^. For all pupil measurements, eyelid closures and blinks were removed and interpolated. Pupil area was normalized between the 1st and 99th percentiles to remove outliers and rescaled to a percentage-based measurement (0-100). Pupil constriction and dilation were classified as the time points where the normalized pupil area was below or above 50% of the maximum value, respectively.

Whisker pad movements were extracted from the average motion energy of the pad. All the extracted behavioral variables were interpolated and upsampled to 20 kHz. To remove noise from the extrema, the average motion energy of the whisker pad was rescaled and normalized as

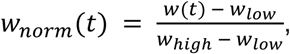

where w(t) is the average motion energy at time t, w_low_ is the 5th percentile, and w_high_ is the 95th percentile. A whisker movement bout was defined as the normalized motion energy surpassing 0.1 and remaining above threshold for at least 0.5 s. Bouts that occurred within 1 s or less were grouped together as a single event.

Running speed was calculated offline by converting analog treadmill voltage (0-3.0 V) to velocity

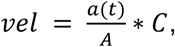

where a(t) is the analog signal at time t, A is the maximum allowed voltage (3.0), and C is the maximum allowed speed (in mm/s). A locomotion epoch was defined by the treadmill velocity reaching 2.5 cm/s and staying above threshold for at least 1 s. Neighboring locomotion bouts that occurred within 1 s were grouped together.

### Biocytin staining and Immunohistochemistry

To obtain histological confirmation of cortical layer for cells recorded in vivo, a subset of neurons were labeled with biocytin during whole-cell recording. Within an hour of experiment termination, animals were deeply anesthetized with isoflurane and perfused through the heart with 1X phosphate-buffered saline (PBS, pH 7.4) followed by 4% paraformaldehyde (PFA) in PBS). Following extraction, the brain was postfixed overnight in 4% PFA at 4°C and transferred to PBS. The tissue was then sectioned into 100 µm thick coronal slices on a vibratome (VT1200, Leica), arranged on a well plate with 200 µL washing solution (PBS 0.1% Triton-X 100 (PBST)) and incubated on a shaker at 37 °C for 2 hours before adding 20 µL of Streptavidin-Alexa Fluor 488 conjugate (S32354, Thermo Fisher, 0.2%). The sections were then shaken and incubated at 37 °C overnight followed by 4 PBST washes every 10 minutes, another round of incubation (2 hours, 37 °C) and 3 more PBST washes every 10 minutes. Lastly, the PBST was replaced with PBS before mounting with an aqueous mounting medium with DAPI (Vectashield). The samples were imaged using a confocal microscope (Zeiss LSM 710) using either a 10X or 20X objective with excitation at 488 nm.

Biocytin staining and immunohistochemistry was combined in a subset of experiments. Following the perfusion and sectioning procedure above, brain slices were washed 4 times in PBS for 10 minutes each. Non-specific antibodies were blocked by a 30 minute wash in 10% goat serum in PBS-T. The sections were then incubated with the primary antibody anti-Ctip2, a marker for cortical neurons with subcerebral projections (rabbit monoclonal, Abcam ab240636, 1:500 final dilution)^82^, 10% goat serum, and 1% bovine serum albumin for 24 hours under gentle agitation at room temperature. After 4 10-minute PBS washes, sections were incubated with gentle agitation at room temperature overnight (12-15 hours) with the conjugated secondary antibody (anti-Rabbit IgG-Alexa Fluor 594, Invitrogen A-11012, 1:500 final dilution), Streptavidin-Alexa Fluor 488 (1:200 dilution), and 2% goat serum. Lastly, the slices were washed 4 times with PBS for 10 minutes each before being mounted. The samples were then imaged as specified above but at 594 nm excitation.

### In vivo plasticity experiments

*Movie clips.* 6 or 10 second clips of continuous scenes from Inception, The Shining, and Fantastic Mr. Fox were used for plasticity experiments. For each experiment, one movie was selected for plateau pairing, and a second movie served as a control (no plateau pairing). Movies were played in a pseudorandom order, with no more than 3 repetitions of the same movie in a row. Approximately 15 repetitions of each movie were played before plateau pairing was performed. After baseline responses to each movie were collected, the bridge balance was checked for full compensation, and a current injection of 800 pA was applied to drive a plateau during the movie selected for pairing. The time of current injection for pairing was chosen by the experimenter and biased towards movie epochs with lower firing rates when possible. 11 of 14 cells received a single, 1 second-long current injection, and the other 3 cells included in the dataset received the following pairings: 20 x 200ms, 10 x 300ms, and 5 x 500ms, noted in Extended Data Fig. 5. Following induction, both movies were played again for at least 15 trials each. For AP pairing, a 1 second-long train of 60 Hz APs was generated by alternating positive (∼1.0 nA) and negative (∼100 pA) 2 ms current injections. The applied hyperpolarizing steps prevented bursting.

#### Static gratings

In a separate cohort of animals, full-field static gratings (as described above in “Visual stimulation”) were used for plasticity experiments. Firing rates during visual stimulation were baseline-normalized, and the evoked spike rate to each orientation was calculated. For each neuron, a non-preferred, non-orthogonal orientation was selected for pairing with a 300 ms-long current injection of 600-800 pA, delayed by 100 ms after stimulus onset. Following pairing, full-field static gratings were presented again.

### In vivo plasticity analysis

#### Movie clips

Voltage signals were processed as described above and separated into trials corresponding to each movie presentation. Trials were aligned either to the start of movie presentation or, for spontaneous plateaus that occurred during grey screen presentation, to the start of the grey screen. Analysis was restricted to a 4-second window (±2 seconds) centered on the leading edge of the plateau or current injection. Action potential-clipped voltage signals were low-pass filtered (<3 Hz) using an FIR filter with a 200 ms Hamming window^20^, then binned into 200 bins.

To identify statistically significant voltage changes, we used a resampling approach to generate 95% confidence intervals (Extended Data Fig. 4). The confidence interval of voltage changes was estimated by randomly assigning trials to “pre” and “post” designations and taking the difference of the averages, 5,000 times. Voltage changes exceeding the 95% confidence interval were considered significant only if they also persisted for at least 0.16 seconds, which was the upper bound of false positive peak durations. We established this duration criterion through 500 simulations across 5 recordings that contained no plateaus but had at least 20 trials each, using randomly selected 4-second analysis windows.

We applied this resampling method to test if significant subthreshold voltage changes manifested in recordings with plateaus or current injections. Only spontaneous plateaus with at least 10 trials of the same movie both before and after their occurrence were included. When assessing voltage changes, if multiple peaks within the 4-second window met both the confidence interval and duration criteria, we selected the peak with the largest magnitude. The timepoint of this peak change in V_s_ was used to report “before/pre” and “after/post” values (see Extended Data Fig. 4 for examples).

To assess firing rate changes, we calculated average firing rates in 100 ms bins before and after the plateau or current injection and then interpolated these to 200 bins to correspond with the subthreshold membrane potential analysis. Firing rates were compared at the timepoint of peak firing rate change identified within the epoch showing significant voltage change.

If a recording had a plateau that reached significance criteria, other plateaus from the same recording were excluded from the analysis. In 3 of 7 recordings with plateaus meeting significance criteria, a second plateau occurred within 300 ms of the first plateau’s end; for these cases, we report the summed duration of both plateaus in Figure 2.

Pairing experiments also included control (unpaired) movies. For control experiments, we compared voltage (or firing rate) at the same timepoint identified for peak change in the paired movie condition.

To examine voltage changes over time, we identified the within-trial timepoint of peak voltage change, then extracted voltage at that timepoint across all trials. These values were baseline-corrected by subtracting the mean voltage from trials before the plateau or current injection. For the plot in Fig. 2d, the number of neurons per time bin are: time (t) = -9, n = 2; t = - 7, n = 2; t = -5, n = 4; t = -3, n = 7; t = -1, n = 7; t = 1, n = 7; t = 3, n = 7; t = 5, n = 5; t = 7, n = 2; t = 9, n = 2; t = 11, n = 2. For Fig. 3d, the number of neurons per time bin are: t = -5.5, n = 6; t = - 4.5, n = 12; t = -3.5, n = 12; t = -2.5, n = 14; t = -1.5, n = 14; t = -0.5, n = 14; t = 0.5, n = 14; t = 1.5, n = 14; t = 2.5, n = 14; t = 3.5, n = 14; t = 4.5, n = 12; t = 5.5, n = 10; t = 6.5, n = 3; t = 7.5, n = 3; t = 8.5, n = 2. For Extended Fig. 7d, the number of neurons per time bin are: t = -4.5, n = 6; t = -2.5, n = 6; t = -0.75, n = 6; t = 0.25, n = 6; t = 1.5, n = 6; t = 3.5, n = 6; t = 5.5, n = 6; t = 7.5, n = 5; t = 9.5, n = 5.

For action potential induction experiments, no significant voltage changes were detected in time windows surrounding the AP train. The voltage change at the timepoint of the first action potential for the population is reported.

#### Static gratings

Firing rates during visual stimulation were baseline-normalized by dividing the evoked spike rate (20 to 200 ms relative to stimulus onset) by the baseline spike rate (-50 to 20 ms). To calculate orientation preference before and after induction, we computed the pre and post global orientation selectivity index (gOSI) as follows:

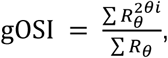

where R is a vector with the average response to each angle θ.

### In vitro electrophysiology

Acute slices were prepared from 8-14 week-old male and female C57/BL6 mice. Sucrose-containing artificial cerebrospinal fluid (sACSF) was used during the slicing procedure, containing (in mM): 90 sucrose, 60 sodium chloride, 26.5 sodium bicarbonate, 2.75 potassium chloride, 1.25 sodium phosphate, 9 glucose, 1.1 calcium chloride, and 4.1 magnesium chloride, with an osmolality of 295-302. The sucrose solution was partially frozen to create a small amount of slush and was kept ice-cold. Artificial cerebrospinal fluid (ACSF) was used for recovery and recording, containing (in mM): 122 sodium chloride, 25 sodium bicarbonate, 3 potassium chloride, 1.25 sodium phosphate, 12 glucose, 1.2 calcium chloride, 1.2 magnesium chloride, 1 ascorbate, and 3 sodium pyruvate, with an osmolality of 302-307. All solutions were saturated with carbogen (5% CO_2_ and 95% O_2_). Mice were anesthetized with isoflurane and decapitated. The brain was extracted in ice-cold sACSF in less than one minute. The brain was blocked at a moderate angle (approximately 20 degrees from coronal) to favor the preservation of L5 apical dendrites in the occipital cortex. After the brain was mounted and submerged in ice-cold sACSF, a vibratome (Leica VT1200S) was used to cut 300 µm-thick slices, which were transferred to ACSF for 30-50 minutes at 36°C. Following recovery, slices were kept at room temperature.

Recordings were performed in ACSF at 34–36 °C. Cells were visualized with an Olympus BX-61 microscope with a water-immersion lens (60X, 0.9 NA; Olympus). L5 PNs were characterized by their large somas 480-630 microns below the pia. Whole-cell current-clamp recordings were obtained with a Dagan BVC-700A amplifier. Patch pipettes with thin-wall glass (1.5/1.0mm OD/ID, WPI) and resistances of 4-7 MΩ were used. Intracellular recording solution contained (in mM): 134 potassium gluconate, 6 KCl, 10 HEPES, 4 NaCl, 4 Mg_2_ATP, 0.3 NaGTP, and 14 phosphocreatine di(tris). Pipette capacitance was fully neutralized prior to break-in, and series resistance was fully bridge balanced throughout the recording, ranging from 10-30 MΩ. The liquid junction potential was not corrected. Current signals were digitized at 20 kHz and filtered at 10 kHz (Prairie View).

### In vitro plasticity experiments

EPSPs were evoked by electrically stimulating axons in layer 5 at ∼100-200 µm lateral from the soma with a bipolar stimulating electrode (ISO-Flex, AMPI) housed in theta glass (2.0/1.4 mm OD/ID). Stimulation intensity was calibrated to generate a small EPSP of 1-3 mV, requiring 5-20 µA of current from the stimulating electrode. Ten minutes of baseline stimulation at 0.1 Hz was recorded to ensure stability of a pair of EPSP pulses (10-20 Hz). A -50 pA hyperpolarizing step was included to estimate input resistance. For pairing, a train of EPSPs (10-20 Hz) was paired with a 300 ms 800 pA current injection at the soma at variable intervals indicated in the text. The timing interval is defined as the time from the middle of the train to the start of the current injection, or vice versa, as previously described^20^. Pairings were repeated 5 times with a 15 second interval between pairing epochs. After the pairing period, EPSPs were monitored for at least 30 minutes. Inclusion criteria were as follows: resting membrane potential could fluctuate from baseline by no more than 3 mV, input resistance could not increase more than 20% of baseline; and series resistance had to remain compensated fully throughout and below 30 MΩ. Cells exhibiting gradual, consistent increases or decreases in EPSP amplitude during the baseline period were excluded. Gabazine (1 µM) was present in ACSF throughout recording. Control experiments were performed as described above, with some modifications. Pre- and post-alone protocols consisted of either only presynaptic stimulation or current injection, respectively. D-AP5 (50 µM) was used for acute blockade of NMDARs and was bath-applied throughout the recording. Experiments with action potential induction consisted of a 60 Hz train of 2 ms 1.0 nA current injections for 300 ms. Hyperpolarizing steps were added between depolarizing steps to prevent bursting during action potential generation. Action potential pairings were repeated 5-10 times to ensure robustness. Layer 4 synapses were tested by placing the stimulating pipette 150 µm above the soma and 50-100 µm away from the apical trunk. All controls with pre- and post-synaptic pairing were performed at the overlapping timepoint (t = 0).

### Statistics

No statistical methods were used to predetermine sample sizes. Sample sizes are similar to those reported in previous publications with related techniques^19,20^. Data were analyzed using two-tailed paired or unpaired non-parametric tests, as stated in the figure legends. Boxplots indicate the 25th and 75th quartiles, with whiskers extending to ±2.7σ. Statistics for Fig. 4 & Supp a-h are as follows: All timepoints vs pre/post only, Kruskal-Wallis test, p = 4.80e-6. 60 Hz, NMDA blockade, & L4 vs pre/post only, Kruskal-Wallis test, p = 0.1731. Means ± SEM and Bonferonni-corrected p-value from Mann-Whitney U-test, comparing the timepoint listed to pre & post only: 0 ms: 1.6643 ± 0.13, p = 2.23e-5. -1250 ms: 1.3566 ± 0.06, p = 3.83e-5. -2250 ms: 1.2820 ± 0.03, p = 6.61e-5. +550 ms: 1.2591 ± 0.06, p = 6.61e-5. -550 ms: 1.3894 ± 0.05, p = 6.61e-5. Means ± SEM for non-significant groups: Post only, 0.9427+-0.01. Pre only, 1.0055 ± 0.01. 60hz, 0.9381 ± 0.02. AP5, 0.9063 ± 0.01. L4, 0.9347 ± 0.02.

## Data Availability

The data that support the findings of this study are available from the corresponding author upon reasonable request.

## Code Availability

Analysis code is available on Github (https://github.com/harnett/neocortical_btsp).

## Acknowledgements

We thank Vincent D. Tang and members of the Harnett laboratory for their comments and suggestions. This work was funded by a Life Science Research Fellowship postdoctoral fellowship (to C.E.Y.), an Emilio Bizzi Fellowship, MathWorks Fellowship, and a Friends of the McGovern Institute fellowship (to R.M.), and a Vallee Foundation Scholars Award and the National Institutes of Health (R01NS106031, R01NS113079, and R01MH135141, to M.T.H).

## Author contributions

C.E.Y., R.M., and M.T.H. designed the experiments. C.E.Y. and R.M. performed in vivo recordings, and C.E.Y. performed in vitro recordings. W.L. and A.B. performed headplate implantation, animal habituation, and histology. W.L. performed immunohistochemistry. A.B. processed behavioral data. C.E.Y. and R.M. analyzed the data and prepared the figures. C.E.Y., R.M., and M.T.H. wrote the manuscript. M.T.H. supervised all aspects of the project.

## EXTENDED DATA

**Extended Data Figure 1.**
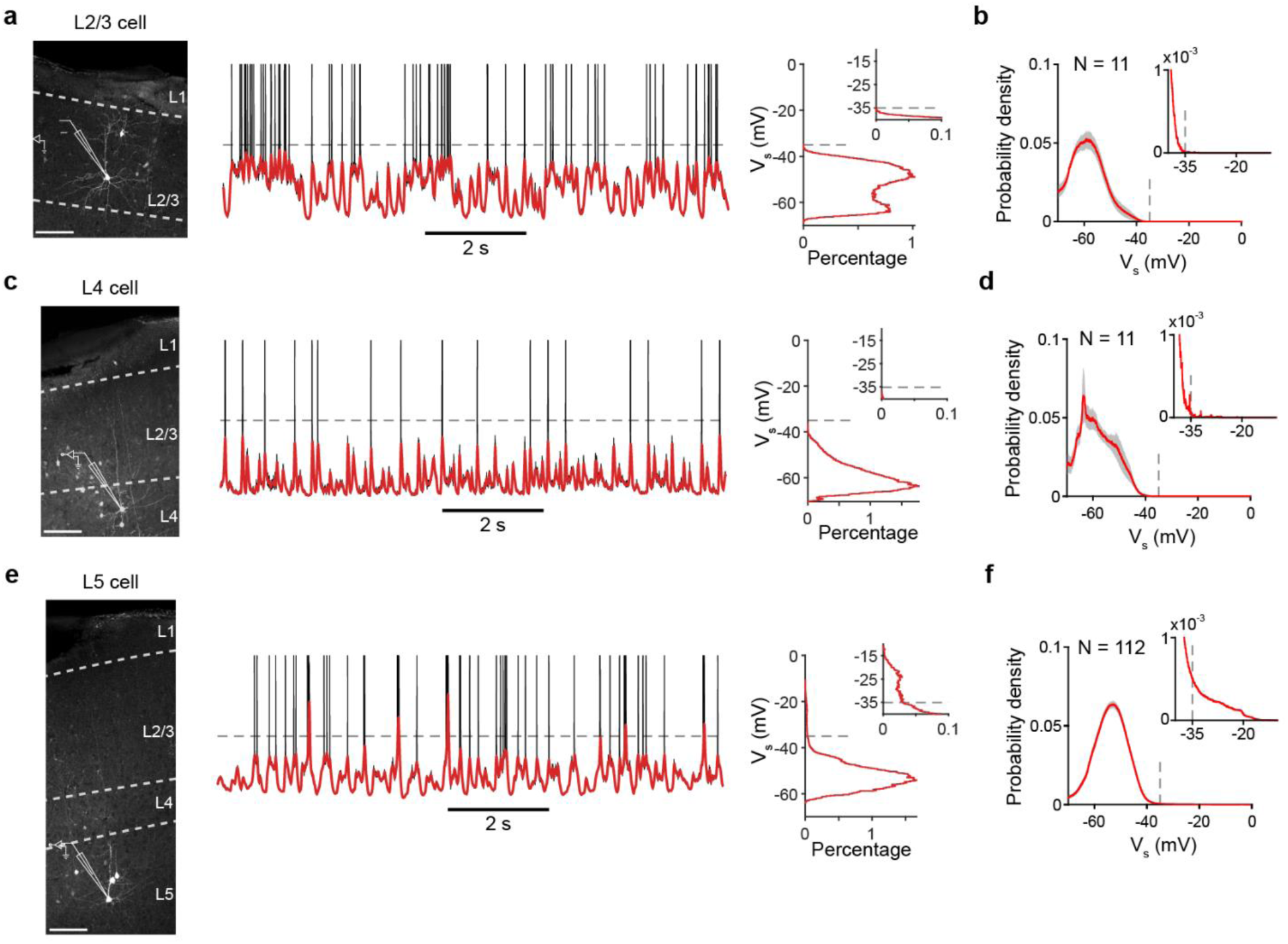
Lack of plateau potentials V1 L2/3 and L4 neurons. **a)** Left, confocal image of an example L2/3 PN that was filled with biocytin during in vivo recording. Scale bar: 100 µm. Middle, representative V_m_ trace (black) and V_s_ (red) for the cell at left. V_m_ traces are clipped at 0 mV for clarity. Right, distribution of filtered V_s_. Inset, distribution tail. **b)** Filtered V_s_ distribution for all sampled L2/3 PNs (n=11). Shaded gray area: S.E.M. Inset: population distribution tail. **c)** As in a, but for a representative L4 neuron. **d)** As in b, for all sampled L4 cells (n=11). **e)** As in a, but for a representative L5 PN. **f)** As in b, for all sampled L5 PNs (n=112). Note the pronounced tail in the inset distribution for L5 from plateau potentials (panel f), which is lacking in L2/3 (panel b) and L4 (panel d).

**Extended Data Figure 2.**
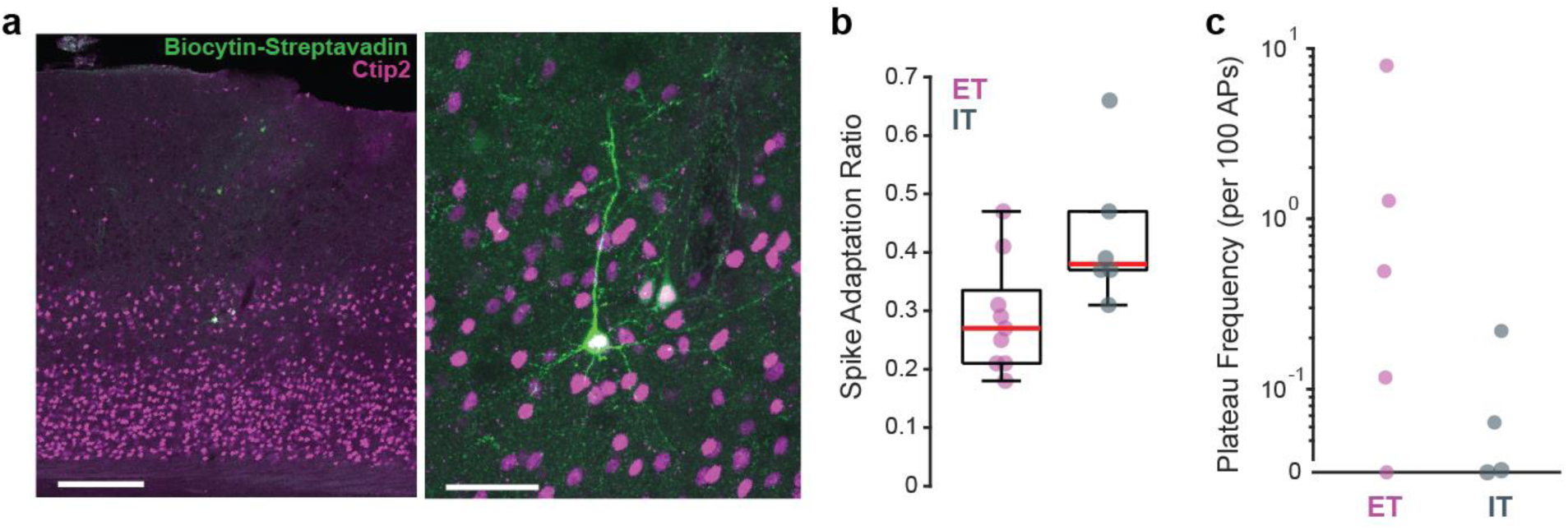
Plateau potentials occur in both ET and IT V1 L5 PNs. **a)** Left, post-hoc biocytin and anti-Ctip2 staining from a L5 PN patch-clamped in vivo. Scale bar: 200 µm. Right, magnified image showing co-localization of the biocytin fill and anti-Ctip2. Scale bar: 50 µm. **b)** In vivo spike adaptation ratio from ET and IT neurons (confirmed posthoc via presence or absence of Ctip2, respectively). Mann Whitney U-test, p = 0.03. Note the overlap in distributions, which prevents clear classification without immunohistochemical confirmation. **c)** Plateau frequency in ET and IT neurons. Mann Whitney U-test, p = 0.17.

**Extended Data Figure 3.**
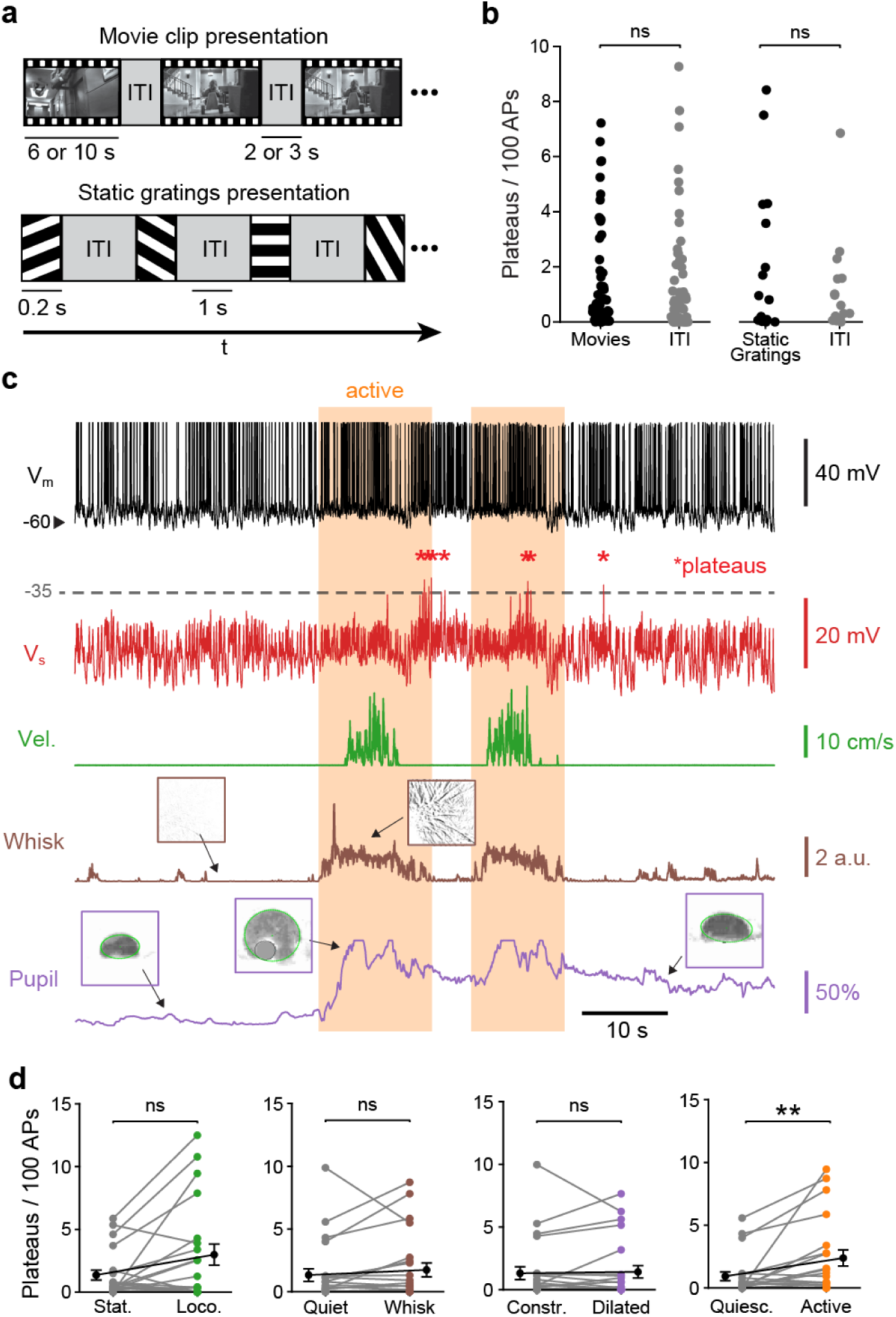
Plateau potential prevalence is not dependent on visual stimulation or behavioral variables. **a)** Schematic of trial structure for movie clip presentations (top) and static grating presentations (bottom). **b)** Plateaus per 100 spikes for movie clips (left, n = 83 L5 PNs) and grating presentations (right, n = 34 L5 PNs). Wilcoxon signed-rank tests, movies vs. ITI, p = 0.5740 (0.97 ± 0.19 and 1.01 ± 0.20); gratings vs. ITI, p = 0.3894 (1.00 ± 0.37 and 0.54 ± 0.22). **c)** V_m_ (black), smoothed V_m_ (V_s_, red), locomotion (green), whisking (brown), and pupil area (purple) from a representative recording. Red asterisks denote plateau potentials. Inset images indicate whisker pad motion energy (top, brown border) during a quiet (left) and an active whisking period (right), and pupil area examples (bottom, purple border). Active periods are indicated in orange. **d)** Plateau probabilities for each binarized state. Wilcoxon signed-rank tests, stationary vs. locomotion, p = 0.1005 (1.36 ± 0.40 and 2.98 ± 0.84); quiet vs. whisking, p = 0.1930 (1.33 ± 0.50 and 1.73 ± 0.55); constricted vs. dilated, p = 0.6051 (1.31 ± 0.51 and 1.42 ± 0.49); quiescent vs. active, p = 0.0065 (0.92 ± 0.35 and 2.38 ± 0.64). Data are mean ± SEM.

**Extended Data Figure 4.**
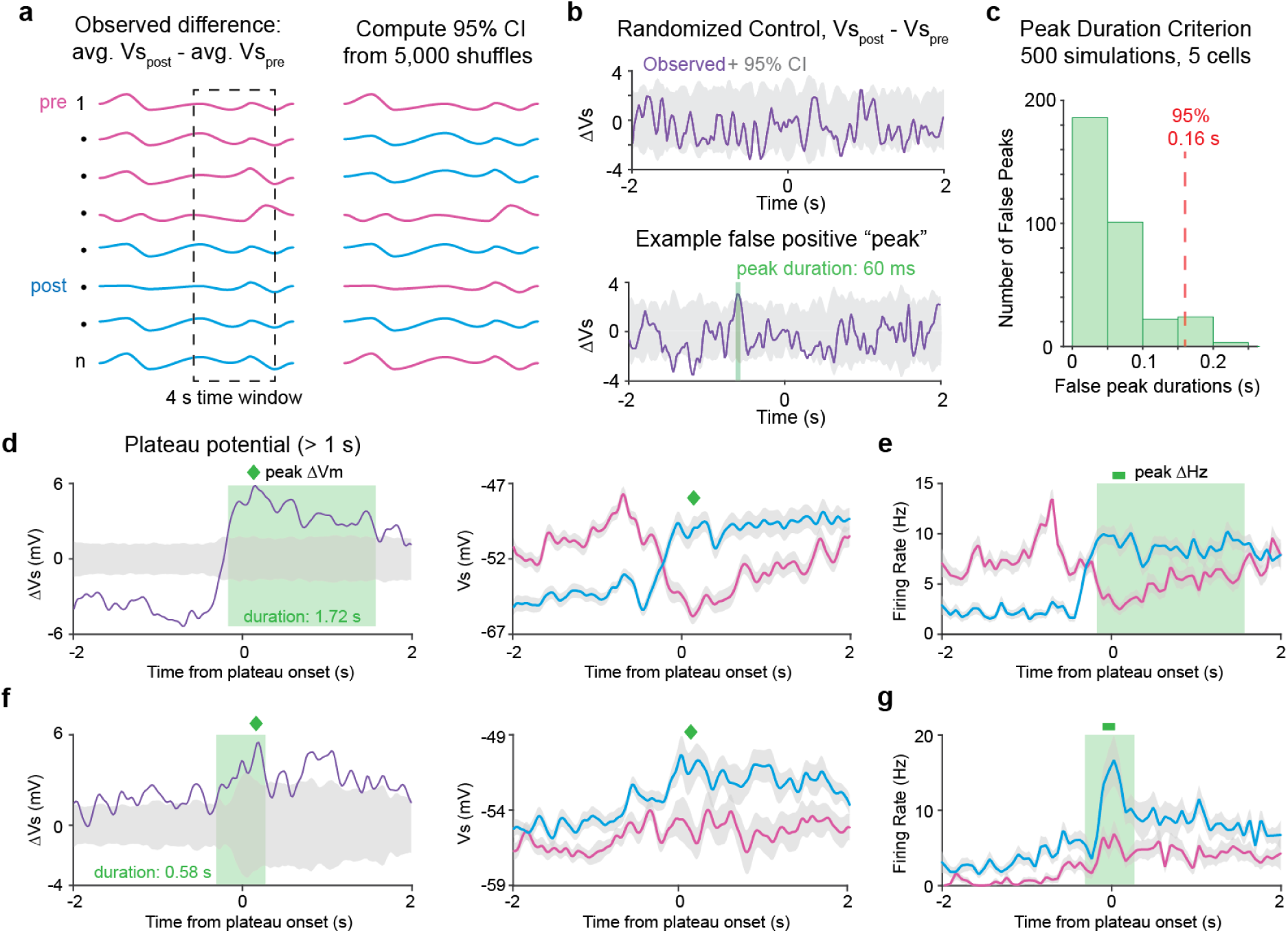
Analysis approach for detection of significant changes in subthreshold membrane potential. **a)** Schematic of trial analysis. Left, pre and post (i.e. before and after) plateau trials are averaged in a 4 second-long window centered around either a spontaneous plateau or a current injection. The difference of these averages is the observed difference (ΔV_s_). Right, the confidence interval of the ΔV_s_ is computed by 5,000 random shuffles of pre and post trials. **b)** Top, example ΔV_s_ across a time window with no plateaus or current injections. Bottom, example false positive peak (green) from another similar control period. **c)** Distribution of false positive peak durations from 500 simulations in 5 cells. Duration criterion is set to peaks > 160 ms. **d)** Left, ΔV_s_ following a long plateau potential (from the cell in Fig. 2a). Note the long duration of the significant epoch (green). The timepoint of peak change is indicated (green diamond) Right, average V_s_ before (pink) and after (blue) the long duration plateau. The timepoint of the peak change (from ΔV_s_ analysis at left) is noted (green diamond) and is used for pre-post comparison. **e)** as in d, but for average firing rate. **f)** and **g)** As in d and e, but for the cell shown in Fig. 2c.

**Extended Data Figure 5.**
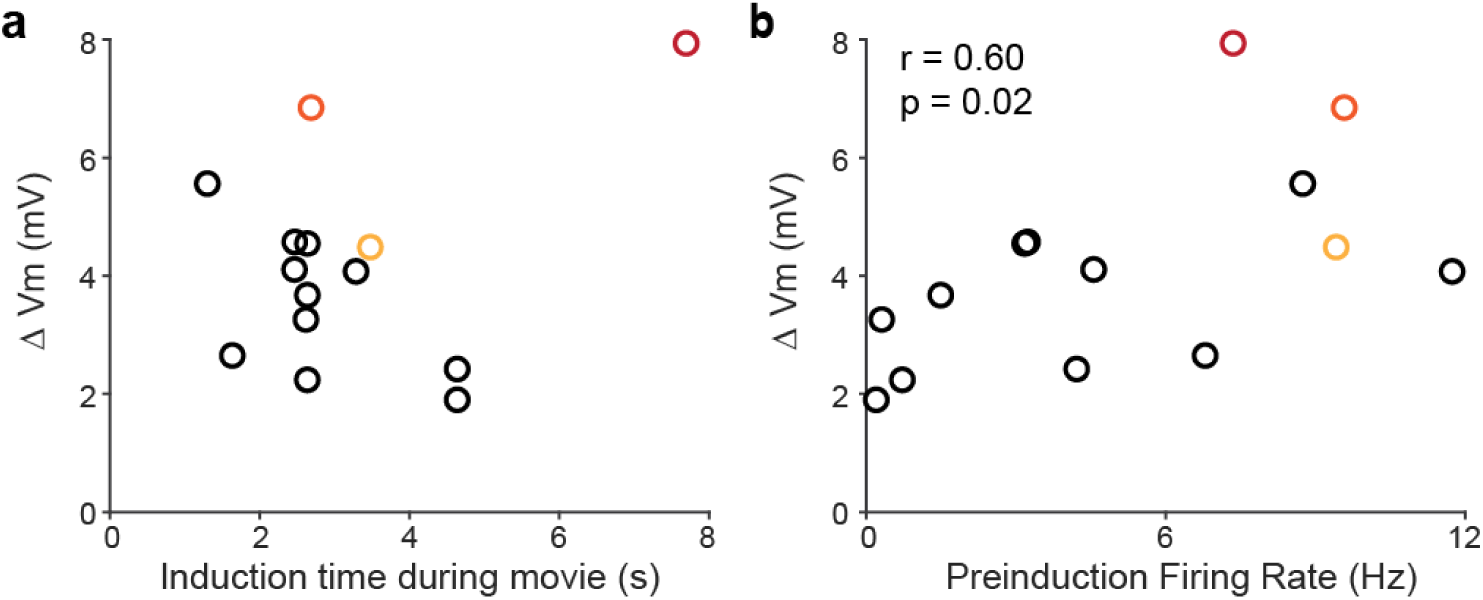
Plasticity as a function of the time of induction during movie presentation and preinduction firing rate. **a)** Average ΔV_s_ of individual L5 PNs as a function of the time of induction (current injection) within a movie. Red, orange, and yellow markers reflect the following induction protocols: 20x200 ms, 10x300 ms, and 5x500 ms, respectively. The 1x1 s protocol was used for all other points (black). Spearman’s correlation, r = -0.04, p = 0.89. **b)** Average ΔV_s_ as a function of neurons’ preinduction firing rate at the same timepoint. Point color coding as in a. Spearman’s correlation, r = 0.60, p = 0.02.

**Extended Data Figure 6.**
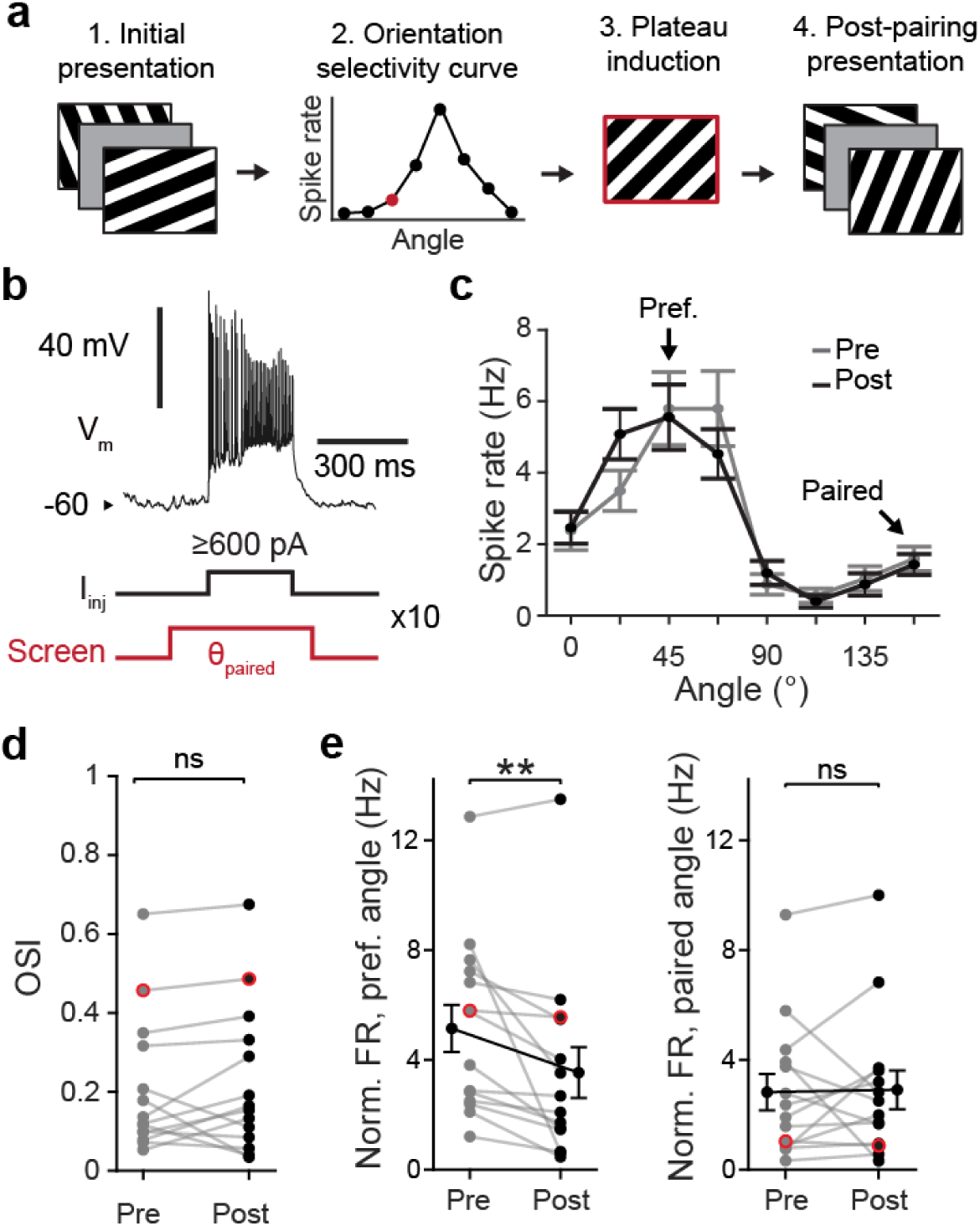
Pairing current injection-evoked plateau potentials with static gratings does not alter orientation selectivity in V1 L5 PNs. **a)** Schematic of experimental design. After 30x 500 ms presentations of all 8 orientations, orientation selectivity curves were built and from this, a non-orthogonal, non-preferred orientation was selected for pairing. After pairing, all 8 orientations were presented again 30x. **b)** Plateau induction protocol. Specific grating orientations (red) were paired with somatic current injection (≥600 pA, 300 ms, black) to evoke plateau-like depolarization, ten times. **c)** Example orientation selectivity curve for a V1 L5 PN before (grey) and after (black) pairing. **d)** Population orientation selectivity indices for n=14 L5 PNs before and after pairing. Example cell from panel c is plotted in red. Paired t-test, pre vs. post, p = 0.4898 (0.21 ± 0.05 and 0.22 ± 0.05). **e)** Left, normalized spike rate at preferred angle before and after pairing. Right, normalized spike rate for paired angles. Example cell from panel c is plotted in red. Wilcoxon signed-rank tests, preferred angle FR pre vs. post p = 0.0013 (5.14 ± 0.86 and 3.53 ± 0.92); paired angle FR pre vs. post p = 0.7032 (2.82 ± 0.66 and 2.91 ± 0.71). Data are mean ± SEM.

**Extended Data Figure 7.**
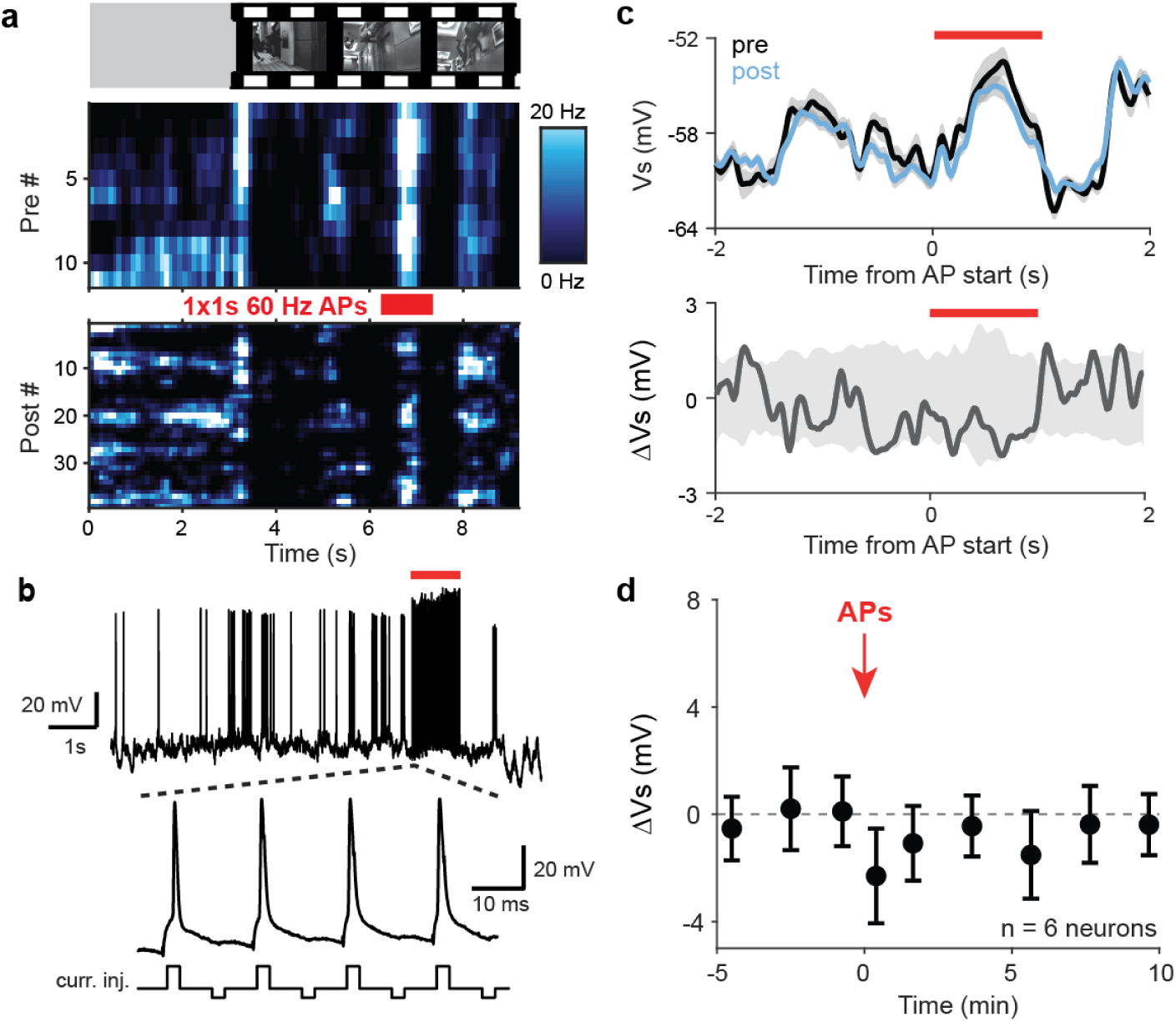
High frequency action potentials without an underlying plateau do not alter subthreshold responses to movie clips. **a)** Firing rate heat maps from a representative neuron pre and post AP pairing. **b)** Intracellular V_m_ during AP pairing. Current injections evoked action potentials at 60 Hz with 2 ms pulses (alternating +1.0 & -0.1 nA,) for a total duration of 1 second. **c)** Top, average V_s_ before and after AP pairing, and bottom, ΔV_s_ with resampled confidence interval (grey). No significant changes in V_s_ were detected. Red bar indicates the start and duration of the action potential train. **d)** Average V_s_ as a function of time for n = 6 neurons used for AP pairing. See Methods for number of neurons in each data point. Data are mean ± SEM.

**Extended Data Figure 8.**
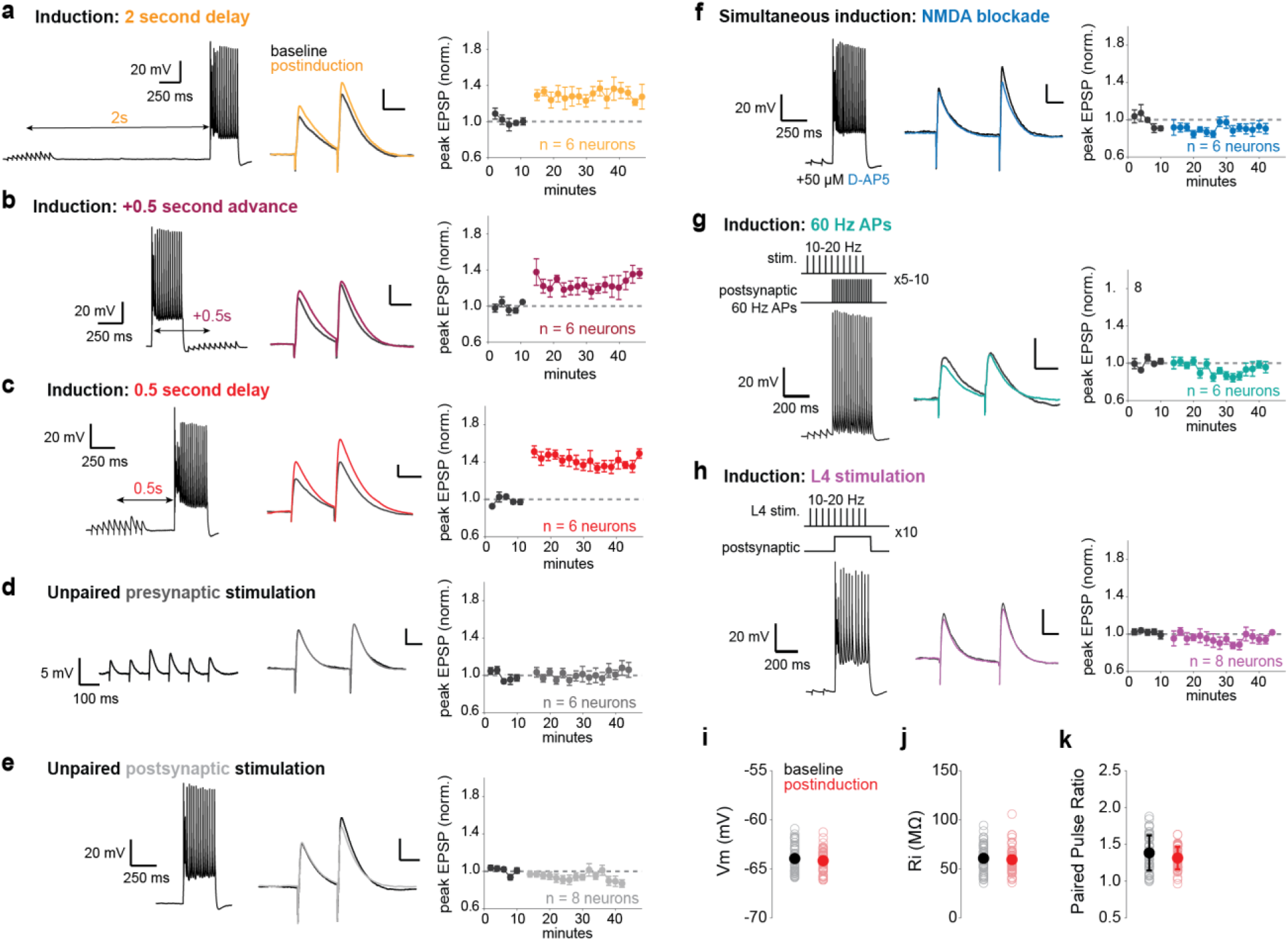
Additional timepoints and controls for ex vivo plasticity experiments. **a)** Left, representative voltage trace of the t = -2.0 s induction paradigm, in which a train of EPSPs driven by electrical stimulation is paired with a somatic current injection, 5 times. Note the current injection occurs at a 2.0 second delay from the middle of the EPSP train. Middle, average paired pulse response during the baseline period (black trace) and the average response postinduction (colored trace). Scale bars: 1 mV, 25 ms. Right, normalized EPSP amplitude for a population of neurons that received the -2.0 s induction paradigm. Induction occurs after baseline collection (black dots), after which the postinduction response is indicated (colored dots). Data are mean ± SEM. **b-h)** As in a, but for other pairing protocols as indicated. See Methods for statistical tests. **i)** Average resting membrane potential for all neurons (n = 67) during baseline (black) and after pairing (colored). Mann Whitney U-test, p = 0.61. Open circles are individual neurons, population mean indicated by filled circles. **j)** As in i, for input resistance. Mann Whitney U-test, p = 0.57. **k)** As in i, for paired pulse ratio. Mann Whitney U-test, p = 0.17.

